# The interaction of physical structure and nutrient loading drives ecosystem change in a large tropical lake over 40 years

**DOI:** 10.1101/2020.10.21.347393

**Authors:** Jemma M. Fadum, Ed K. Hall

## Abstract

Many lakes across the world are entering novel states and experiencing altered biogeochemical cycling due to local anthropogenic stressors. In the tropics, understanding the drivers of these changes can be difficult due to a lack of documented historic conditions or an absence of continuous monitoring that can distinguish between intra- and inter-annual variation. Over the last forty years (1980-2020), Lake Yojoa (Honduras) has experienced increased watershed development as well as the introduction of a large net-pen Tilapia farm, resulting in a dramatic reduction in seasonal water clarity, increased trophic state and altered nutrient dynamics, shifting Lake Yojoa from an oligotrophic (low productivity) to mesotrophic (moderate productivity) ecosystem. To assess the changes that have occurred in Lake Yojoa as well as putative drivers for those changes, we compared Secchi depth (water clarity), dissolved inorganic nitrogen (DIN), and total phosphorus (TP) concentrations at continuous semi-monthly intervals for the three years between 1979-1983 and again at continuous 16-day intervals for 2018-2020. Between those two periods we observed the loss of a clear water phase that previously occurred in the months when the water column was fully mixed. Seasonal peaks in DIN coincident with mixing suggest that an enhanced accumulation of ammonium in the hypolimnion (the bottom layer of a stratified lake) during stratification, and release to the epilimnion (the top layer of a stratified lake) with mixing maintains high algal abundance and subsequently low secchi depth during what was previously the clear water phase. This interaction of nutrient loading and Lake Yojoa’s monomictic stratification regime illustrates a key phenomenon in how physical water column structure and nutrients interact in tropical monomictic lakes. This work highlights the need to consider nutrient dynamics of warm anoxic hypolimnions, not just surface water nutrient concentrations, to understand environmental change in these societally important but understudied ecosystems.

## 1. Introduction

It is increasingly evident that human activities (both local and global) are altering lake ecosystems around the world. The warming of surface waters due to climate change is altering the duration of stratification and water column stability (Pilla et al. 2020) as well as intensifying eutrophication (Moss et al. 2011). In some regions, increased nitrogen supply has shifted the macronutrient that limits primary productivity from nitrogen (N) to phosphorus (P) (Elser et al. 2009) and altered dissolved N:P stoichiometry of lakes (Isles et al. 2017). These impacts are often exacerbated by more local aspects of global change including changes in land use that can increase the loading of nutrients (Carpenter et al. 1998), metals (Yu et al. 2014), and organic pollutants (Lorgeoux et al. 2016) from the surrounding watershed. In-lake activities, such as aquaculture, are also altering key biogeochemical pathways (Christensen et al. 2000).

Understanding how these changes are affecting lakes in the tropics presents a challenge because of the paucity of monitoring efforts in many tropical regions (Bennun 2001; Riveros-Iregui et al. 2018). Without long-term datasets, it is difficult to distinguish between intra-annual variability, periodic or semi-periodic inter-annual variability (e.g., changes associated with El Niño-Southern Oscillation), and directional inter-annual changes. While studies of the African Great Lakes (Talling and Lemoalle 1998; Hecky 2000; Bootsma and Hecky 2003), Central American lakes such as Lake Atitlán (Corman et al. 2015; Weisman et al. 2018) and Brazilian reservoirs and floodplain lakes (Tundisi et al. 1993; Melack et al. 2020) among many others, have made considerable contributions to our understanding of tropical lake ecosystems, time series data remain far less common in tropical compared to temperate watersheds (Riveros-Iregui et al. 2018).

Despite the critical role temperate lakes play in the understanding of aquatic ecosystem biogeochemistry, we cannot directly translate knowledge derived from the long-term monitoring of temperate lakes to tropical lakes because of differences in origin, basin morphology, hydrological drivers, climate, and seasonality that occur across latitudinal gradients. For example, temperature is unlikely to directly limit biogeochemical rate processes in tropical lake ecosystems (Talling and Lemoalle 1998). However, temperature does have an indirect impact on biogeochemical rate process in the tropics by influencing stratification and water column stability (MacIntyre and Melack 2009).

To assess changes in water transparency, nutrient dynamics, shift in trophic state, and potential drivers of these changes in a large monomictic tropical lake, we compared two intensive sampling campaigns in Lake Yojoa that took place forty years apart. The first campaign sampled Lake Yojoa at three locations semi-monthly from 1979 to 1983 (henceforth referred to as ‘historic sampling’, Vaux and Goldman 1984) and provides a rare opportunity to assess the historic conditions of a tropical lake. To create a comparative contemporary dataset, we sampled the lake at five pelagic stations (including three sites aligned with the three historic locations) every 16 days from 2018 to 2020. We also conducted bioassays during mixed (January) and stratified (June) conditions to assess the dominant limiting nutrient to algal productivity. We then compared our contemporary results to those from the historic period. The principal objectives of this study were to 1) identify changes in water transparency and macronutrient (N and P) concentrations over the past forty years and classify the corresponding shift in trophic state, 2) identify differences in inter and intra-annual patterns in these parameters and 3) evaluate the potential drivers of these observed changes to make the findings applicable to other monomictic tropical lake ecosystems.

## 2. Methods

### 2.1 Study site

Lake Yojoa, located in west central Honduras (Figure 1), is the largest freshwater lake in the country (maximum depth = 27.3 m, mean depth= 14.6 m, Supplementary Table 1) and the economic backbone of the surrounding region, supporting local livelihoods (e.g., fishing, agriculture, recreation, tourism, Studer 2007). Today, the Lake Yojoa watershed houses nine municipalities with an estimated combined population of 74,624 (Rivera 2003) after seeing rapid population growth of approximately 68% in recent decades (House 2002). In addition to increasing anthropogenic activity in the watershed, industrial scale net-pen Tilapia production was introduced in 1997. As of 2016, Regal Springs Tilapia operated 106 Tilapia cages in Lake Yojoa (Monterey Bay Aquarium Seafood Watch 2017).

**Figure 1.**
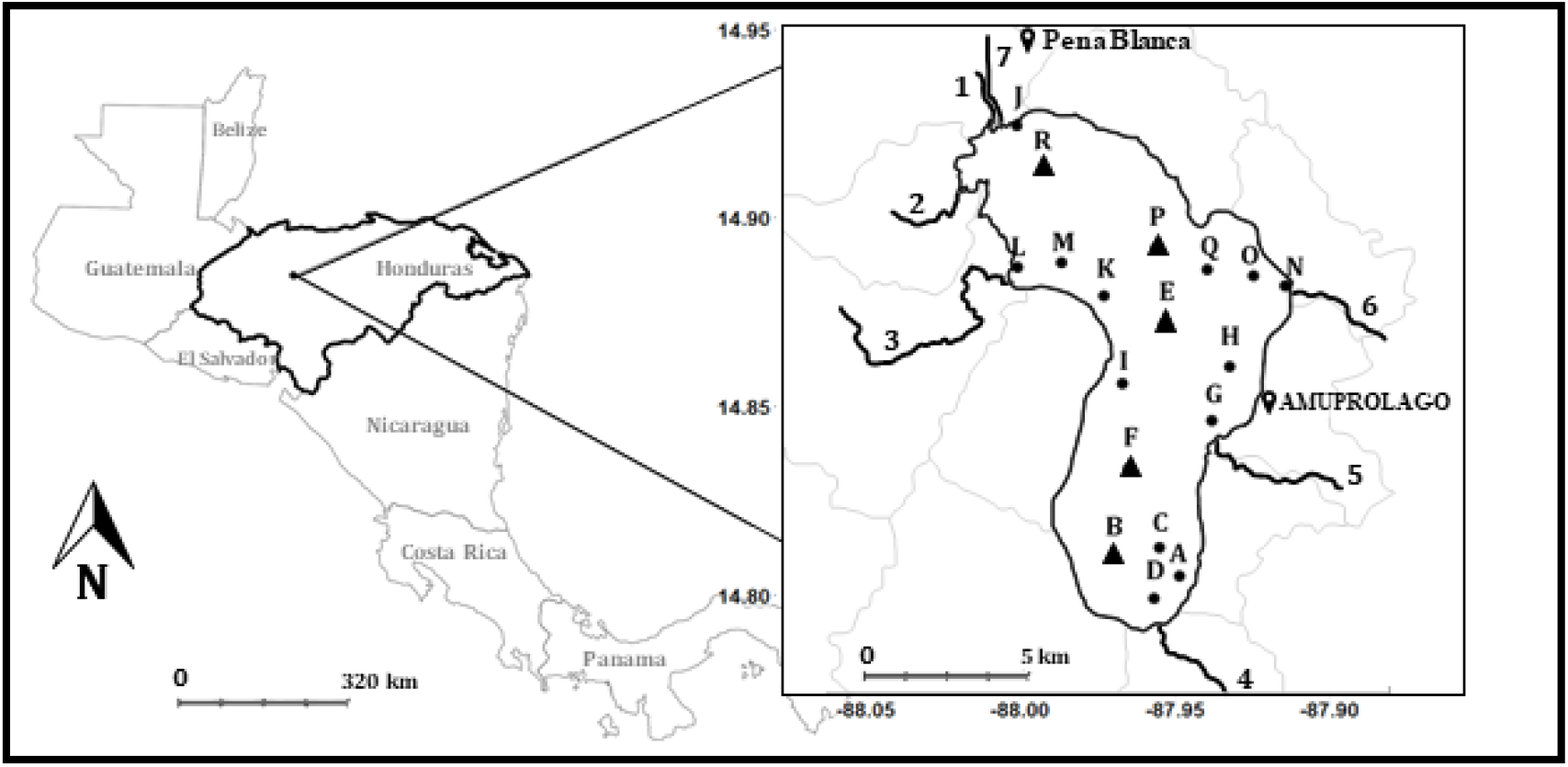
Contemporary sampling stations within Lake Yojoa for twice-annual (A-R) and semi-monthly (B, E, F, R and P) sampling. Principal tributaries 1) Rio Helado 2) Rio Balas 3) Rio Raices 4) Rio Varsovia 5) Rio Cacao 6) Rio Yure and outflow 7) Canal

### 2.2 Sample collection

All historic data are from the El Cajon Project (Vaux and Goldman 1984). From September of 1979 to February of 1983, samples were collected semi-monthly at three pelagic stations. The most consistently sampled location from the historic sampling (“Index”) aligns with contemporary station E, while “Station 2” aligns with contemporary station F, and “Station 1” is between stations R and P (Figure 1). Briefly, samples from both the epilimnion and the hypolimnion were analyzed for total phosphorus (TP), ammonium (NH_4_^+^), and nitrate (NO_3_^-^) as described below. Additionally, on each sampling day, temperature, and dissolved oxygen (DO) profiles of the water column were collected and Secchi depth was also measured (Vaux and Goldman 1984). Full details of sampling and analyses methods are available in Vaux and Goldman (1984).

For the contemporary sampling campaign, in collaboration with the Asociación de Municipios del Lago de Yojoa y su Área de Influencia, AMUPROLAGO, we collected water samples and took Secchi depth measurements and DO and temperature profiles every 16 days from March 2018 to January 2020. In addition to the semi-monthly epilimnetic (surface water) sampling at five stations, in June of 2018 and 2019 and January of 2019 and 2020 we collected epilimnetic and hypolimnetic samples at each of the five regular stations (B, F, E, P and R) and 13 additional stations (18 sites in total, Figure 1). For all sampling efforts, temperature and DO profiles were measured at two-meter intervals using a Hydrolab MS5 Multiparameter Sonde (OTT HydroMet, Loveland, CO). Epilimnetic samples were collected at one meter below the surface with an opaque Van Dorn water sampler and temporarily stored in opaque HDPE Nalgene bottles and placed in a cooler. Hypolimnetic samples were collected at all locations deep enough to be stratified during June sampling events. Samples were collected at a depth of two meters above the sediment (depths between 17-23 m) to ensure sampling occurred below the thermocline (∼12 m) but above the sediment to avoid contamination. Whole water samples were frozen for TP analysis while NH_4_^+^ and NO_3_^-^ samples were first filtered through 25 mm glass microfiber filters, Grade GF/F via Polysulfone filter funnels and subsequently frozen. All samples were stored in 15 ml centrifuge tubes before being transported to Fort Collins, CO, USA while frozen. Analyses were performed at the EcoCore facility at Colorado State University, Fort Collins, CO, USA as described below. Contemporary precipitation data were collected for this study at the AMUPROLAGO office (Figure 1) via HOBO U30 USB Weather Station Data Logger (Part # U30-NRC).

### 2.3 Nutrient enrichment bioassays

In order to directly address whether N or P were primarily limiting algal productivity, during January and June of 2019, forty-eight clear glass bottles (120 mL) were filled with lake water from station E after passing the water through an 80 μm mesh to remove grazers. In addition to the unamended control group, three treatment groups were prepared with twelve replicates each: +N (as NH_4_SO_4_, Sigma-Aldrich cat. no. A4418-100G), +P (as K_2_PO_4_, Sigma-Aldrich, cat. no. P3786-100G) and +NP. Concentrations of nutrient amendments were chosen to approximately double *in situ* concentrations of DIN and TP based on epilimnetic nutrient concentrations from March and June of 2018. N and P amendments in January 2019 were 6.92 µM N and 0.24 µM P, final concentration. In June 2019, N and P amendments were 3.15 µM N and 0.94 µM P, final concentration. We incubated each bottle in an outdoor water bath positioned near the lake as to expose bottles to ambient surface light conditions but buffer diel changes in air temperature. Between the start and end of the experiment bottles were agitated daily to minimize settling of algae. After three days, bottles were shaken gently but thoroughly to homogenize the bottle and algae from each bottle were collected on GF/F filters. Filters were folded, wrapped in aluminum foil, and immediately frozen for transport to Fort Collins, CO. Within three days of sampling, chlorophyll a (Chl-a) was measured on a 10-AU fluorometer (Turner Designs Part No. 10-AU-074, calibrated using Turner Designs liquid Chl-a standards) via extraction in 90% acetone with acidification to 0.003 N HCl with 0.1 N HCl (Arar 1997). Algal response for each treatment was recorded as percent increase from the mean algal concentration within the control treatment. We used pre-acidification values (Chl-a plus phaeophytin) for all controls and treatments to represent total changes in algal biomass.

### 2.4 Nutrient analyses

During the historic study, NH_4_^+^ was measured using the indophenol method (Solórzano 1969), NO_3_^-^ was measured using the hydrazine reduction method (Kamphake et al. 1967), and TP was first digested to soluble reactive phosphorus (SRP) via acid hydrolysis (using H_2_SO_4_) then measured using the ascorbic acid-molybdate method (Murphy and Riley 1962). For the contemporary sampling campaign, TP was first digested to SRP via the Alkaline Potassium Persulfate method under sub-boiling (90 ºC) temperature, then measured using a colorimetric ascorbic acid assay (EPA Method 365.3 in U.S. Environmental Protection Agency 1983) modified to be analyzed on a UV-STAR 96-well Microplate via Infinite M200 TECAN at 880 nm absorbance detection. NH_4_^+^ and NO_3_^-^ were measured using a Flow Solution FS 3700 Automated Chemistry Analyzer (O.I. Analytical, College Station, TX). NH_4_^+^ was determined by an indophenol method (German Standard Methods, DIN in Society of German Chemists 1982). NO_3_^-^ was determined via automated colorimetry (EPA Method 353.2 in O’Dell 1993) which relies on determination via diazotizing with sulfanilamide and coupling with N-(1-naphthyl)-ethylenediamine dihydrochloride and nitrate reduction to nitrite in a cadmium column prior to mixing with the color reagent. NH_4_^+^ and NO_3_^-^ values are presented separately as well as combined and reported as dissolved inorganic nitrogen (DIN).

### 2.5 Calculations and statistical analyses

Data analyses were performed in R (version 4.0.2) using packages tidyverse (Wickham et al. 2019), ggplot2 (Wickham 2016), multcompView (Graves et al. 2019) and agricolae (de Mendiburu 2020) and base R. We calculated Schmidt stability index (SSI) values using rLakeAnalyzer (Winslow et al. 2019). We used one-way analysis of variance (ANOVA) to test for statistically significant differences for intra-annual and inter-annual comparisons of Secchi depth, DIN and TP. Treatment groups in the nutrient enrichment bioassay were compared using Tukey’s HSD (honestly significant difference) test.

To make our results more comparable to other studies on changing lake trophic state, we calculated trophic state index (TSI) for the lake for each sampling campaign using Secchi depth according to Carlson (1977). We chose this method over similar metrics such as Toledo (1983), Salas and Martino (1991), Lamparelli (2004), or Cunha (2013) due to the reliability and consistency of Secchi depth compared to other parameters, absence of Chl-a values from the historic campaign and concerns over under-estimation of trophic state of some models (Klippel et al. 2020). In Lake Yojoa, the majority of variance in Secchi depth can be explained by shifts in algal abundance, making Secchi depth an appropriate tool for estimating trophic state where historic Chl-a values are unavailable (Appendix A).

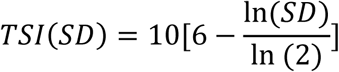

In addition to the nutrient enrichment bioassays, we compared the molar ratio of dissolved inorganic N to Total P (DIN:TP) in the epilimnion between the two sampling periods to use as a proxy for potential macronutrient limitation of primary productivity (Wetzel 2001; Bergström 2010). We used the ratio DIN:TP> 10 to suggest P limitation, 10>DIN:TP>5 to suggest co-limitation and DIN:TP <5 to suggest N limitation (Wetzel 2001).

## 3. Results

### 3.1 Stratification

To assess the mixing regime of contemporary Lake Yojoa, we analyzed DO profiles from sampling events across the two years of contemporary monitoring. For the years of our study, Lake Yojoa behaved principally as a monomictic system, with stratification beginning in February with the thermocline fully establishing at a depth of 12 m in April and water column mixing occurring again in November of each year (Figure 2a). We did observe several partial mixing events during the stratified periods, but these were localized and/or intermittent and stratification was quickly reestablished (Figure 2a). Temperature profiles between the two sampling periods differed, with increased temperature (mean change of 3.0 ºC) in the contemporary relative to the historic period across all depths (Figure 2b). Mean monthly SSI values indicated that water column stability was not significantly different between historic and contemporary sampling periods for nine of the eleven months we were able to compare (Figure 2c).

**Figure 2.**
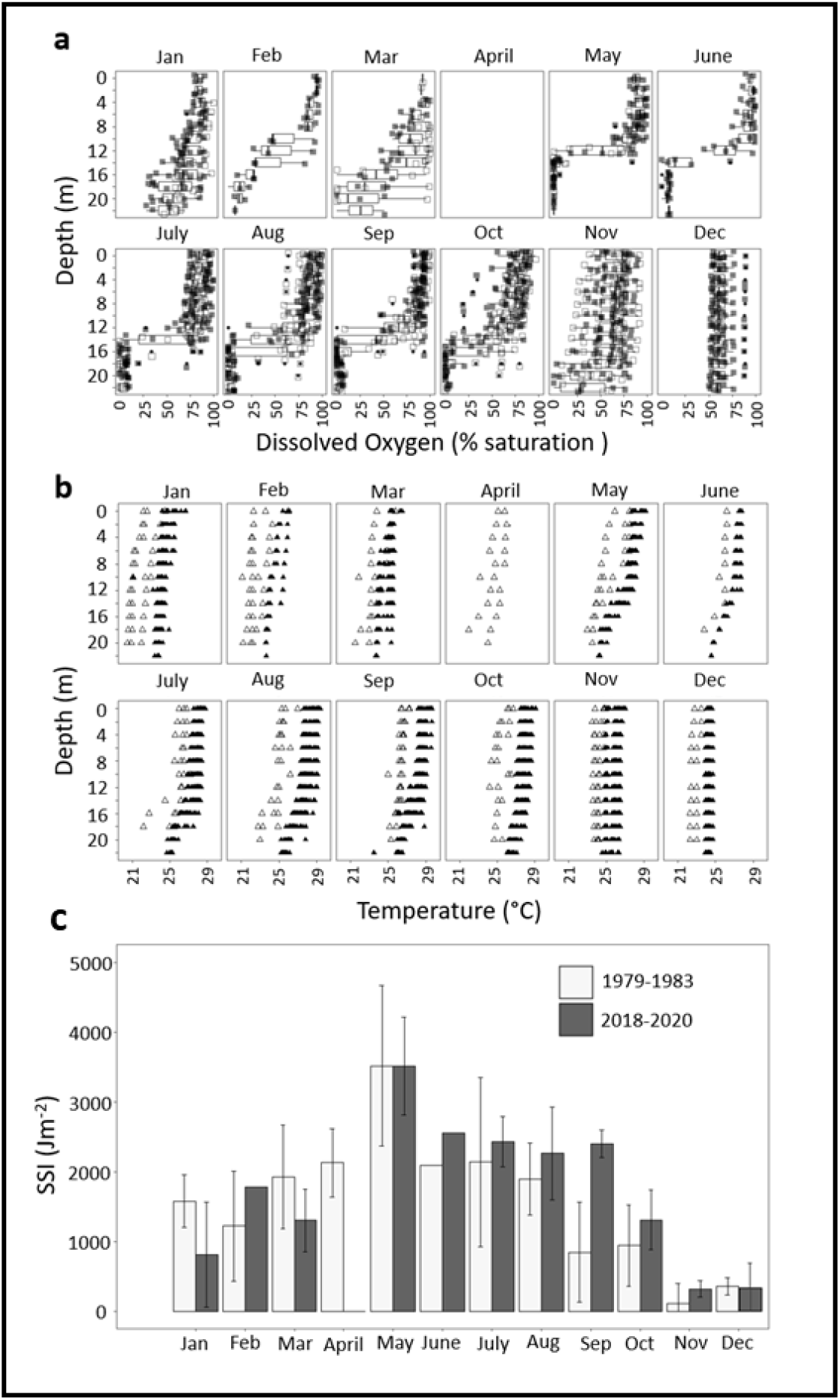
**a)** Dissolved oxygen profiles for contemporary sampling for all months (1-12) in 2019 (open squares) and 2020 (filled squares) over two-year boxplots (min, interquartile range, and max). **b)** Temperature profiles for each month of all years of contemporary (filled triangles) and historic (open triangles) sampling. Due to sonde malfunction and the COVID-19 pandemic, profile data is unavailable for April of both 2019 and 2020. **c)** Monthly mean Schmidt Stability Index (SSI) values ± standard deviation.

### 3.2 Secchi depth and trophic state

We assessed changes in Lake Yojoa’s trophic state between the two sampling periods (inter-annual) and between months within each sampling period (intra-annual) by comparing mean Secchi depths. Mean annual Secchi depth and TSI from the historic period (mean Secchi depth = 7.3 ± standard deviation 2.1 m, and TSI = 31.1 ± 2.7) were significantly different (*p*<0.001, df = 107) than present day conditions (mean Secchi depth = 3.2 ± 1.0 m, and TSI = 43.3 ± 31.5).

The differences in annual mean transparency and trophic state between each sampling period were driven by changes in intra-annual dynamics.

During the historic sampling period, Lake Yojoa experienced a clear water phase from August to March (mean Secchi depth = 8.25 ± 1.7 m and TSI = 29.8 ± 2.7) with annual minimum Secchi depth (mean Secchi depth = 5.0 ± 1.1 m and TSI = 37.0 ± 3.4) occurring between April and July (Figure 3a). However, in the contemporary sampling, we observed no pronounced change in water clarity among seasons. Secchi depths for the months that previously had the minimum secchi depths (April to July) had, on average, 36% lower transparency (mean Secchi depth = 3.2 ± 1.0 m and TSI = 44.2 ± 4.8) in the contemporary sampling. Similarly, the months that previously had a clear water phase (August – March) were statistically indistinguishable (*p*=0.7, df= 87) from the months that had the annual minimum (mean Secchi depth = 3.2 ± 1.0 m and TSI= 43.7 ± 4.1), resulting in a complete loss of seasonality of water clarity between the two sampling periods.

**Figure 3.**
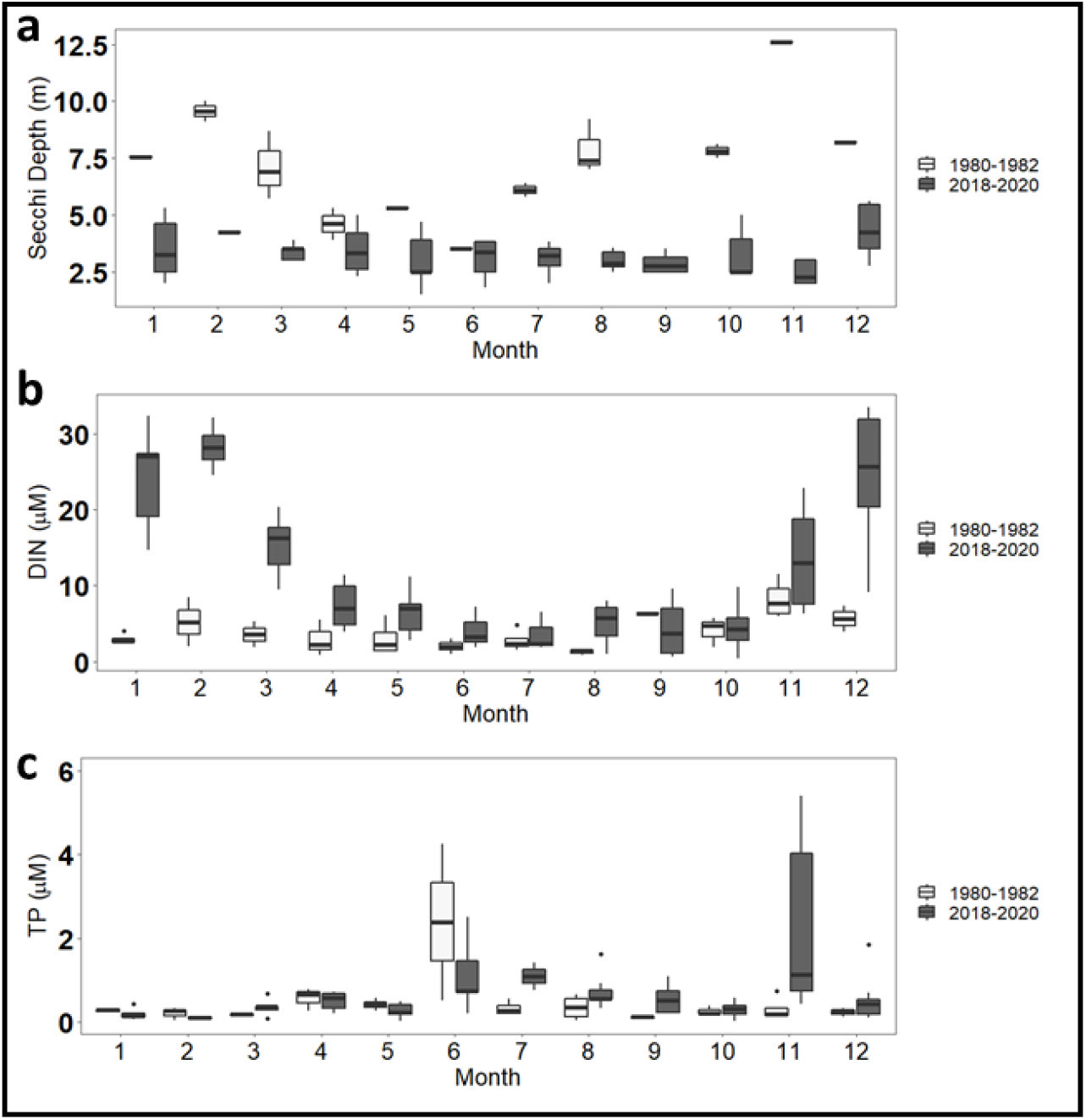
**a)** Secchi depths **b)** dissolved inorganic nitrogen (DIN) and **c)** total phosphorus (TP) at station E. Samples collected from the epilimnion at a depth of 1 meter.

### 3.3 Inorganic nitrogen and total phosphorus

To evaluate putative drivers of changes in the trophic state of Lake Yojoa we also assessed inter-annual and intra-annual changes in epilimnetic nutrient concentrations between contemporary and historic periods. During the historic period, DIN varied throughout the year but had no clear seasonality (mean DIN = 3.99 ± 2.61 μM, Figure 3b). However, during contemporary periods, mean annual DIN and intra-annual variation in DIN were greater than that observed during the historic period (mean DIN= 12.28 ± 9.85 μM, Figure 3b) with greater variance between contemporary and historic periods (DIN variance = 97.1 vs. 6.80, respectively). The increase in contemporary DIN was most pronounced directly following mixing in November (Figure 2a and 3b). After mixing, epilimnetic concentrations of DIN remained significantly higher (*p*<0.001, df=65) than pre-mixing levels until the onset of stratification in April when epilimnetic DIN concentrations returned to pre-mixing levels (Figure 3b).

In contrast to DIN, contemporary epilimnetic TP values did not have increased seasonality in the contemporary period compared to the historic period (Figure 3c). Mean annual TP values increased between historic and contemporary periods (0.41 ± 0.67 μM and 0.72 ± 1.02 μM, respectively), but these differences were not significant (*p*=0.1, df = 117). There was a pronounced increase in epilimnetic TP in June compared to other months of the year that was significant in the historic period (*p*<0.05 for all months) but not significantly different from the other months (*p*>0.5 for all months) in the contemporary sampling. We also observed a secondary peak in the contemporary data in November, coincident with mixing. Contemporary epilimnetic TP in November (2.20 ± 1.93 μM) was somewhat higher relative to the historic period (0.31 ± 0.29 μM), but this difference was only marginally significant (*p*=0.07, df=14, Figure 3c).

We also compared differences in hypolimnetic concentrations of TP, NH_4_^+^ and NO_3_^-^ between historic and contemporary periods during mixed (January) and stratified (June) sampling events to test if the increase in epilimnetic DIN and TP following November mixing was driven by changes in hypolimnetic N and P between sampling periods. During the historic period, there was no significant difference between epilimnetic or hypolimnetic concentrations of NO_3_^-^ (*p*=0.504 df=9), TP (*p*= 0.884, df=9), or NH_4_^+^ (*p*= 0.298, df=9) when the lake was stratified (Table 1). In the contemporary sampling period, we observed no significant (*p*=0.67, df=5) difference in NO_3_^-^ between the two strata (Table 1). However, in the contemporary sampling period, during stratification, the hypolimnion of Lake Yojoa was significantly enriched in NH_4_^+^ (*p*<0.001, df=5) and marginally enriched in TP (*p*= 0.078, df=7) relative to the epilimnion (Table 1), representing ∼7.6 times more NH_4_^+^ and ∼2.3 times more TP than was observed during stratification in the historic data (Table 1). The initial increase in epilimnetic DIN following mixing was driven by an abrupt increase in NH_4_^+^, that was quickly converted to NO_3_^-^ (Figure 4). NO_3_^-^ remained the dominant DIN species in the epilimnion throughout the duration of the mixed water column phase (Figure 4).

**Table 1.**
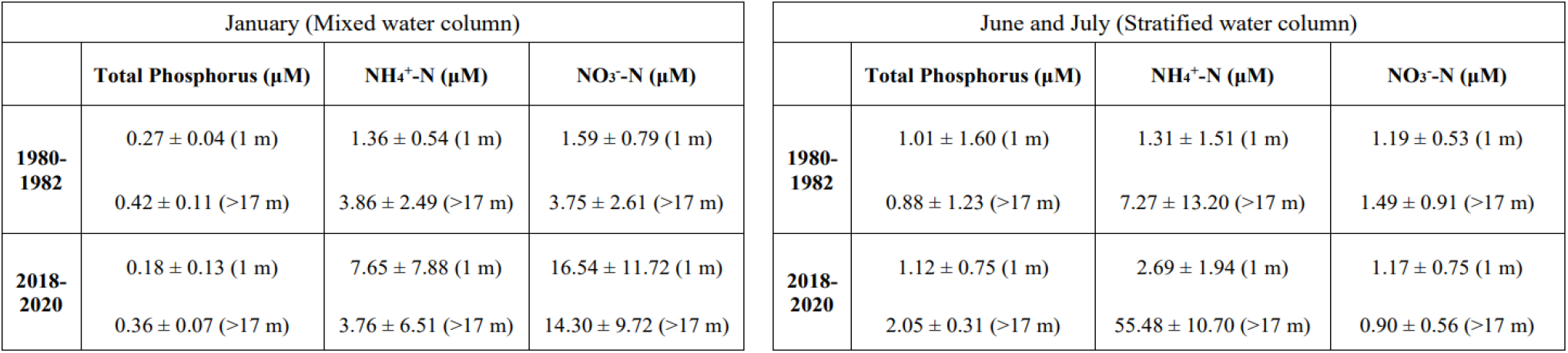
Epilimnetic and hypolimnetic nutrient concentrations for each sampling campaign at station E (mean ± sd). Values are provided next to sampling depth.

| January (Mixed water column) |  |  |  | June and July (Stratified water column) |  |  |  |
| --- | --- | --- | --- | --- | --- | --- | --- |
| | Total Phosphorus ($\mu\text{M}$) | $\text{NH}_4^+\text{-N}$ ($\mu\text{M}$) | $\text{NO}_3^-\text{-N}$ ($\mu\text{M}$) | | Total Phosphorus ($\mu\text{M}$) | $\text{NH}_4^+\text{-N}$ ($\mu\text{M}$) | $\text{NO}_3^-\text{-N}$ ($\mu\text{M}$) |
| **1980-1982** | $0.27 \pm 0.04$ (1 m) | $1.36 \pm 0.54$ (1 m) | $1.59 \pm 0.79$ (1 m) | **1980-1982** | $1.01 \pm 1.60$ (1 m) | $1.31 \pm 1.51$ (1 m) | $1.19 \pm 0.53$ (1 m) |
| | $0.42 \pm 0.11$ (>17 m) | $3.86 \pm 2.49$ (>17 m) | $3.75 \pm 2.61$ (>17 m) | | $0.88 \pm 1.23$ (>17 m) | $7.27 \pm 13.20$ (>17 m) | $1.49 \pm 0.91$ (>17 m) |
| **2018-2020** | $0.18 \pm 0.13$ (1 m) | $7.65 \pm 7.88$ (1 m) | $16.54 \pm 11.72$ (1 m) | **2018-2020** | $1.12 \pm 0.75$ (1 m) | $2.69 \pm 1.94$ (1 m) | $1.17 \pm 0.75$ (1 m) |
| | $0.36 \pm 0.07$ (>17 m) | $3.76 \pm 6.51$ (>17 m) | $14.30 \pm 9.72$ (>17 m) | | $2.05 \pm 0.31$ (>17 m) | $55.48 \pm 10.70$ (>17 m) | $0.90 \pm 0.56$ (>17 m) |

**Figure 4.**
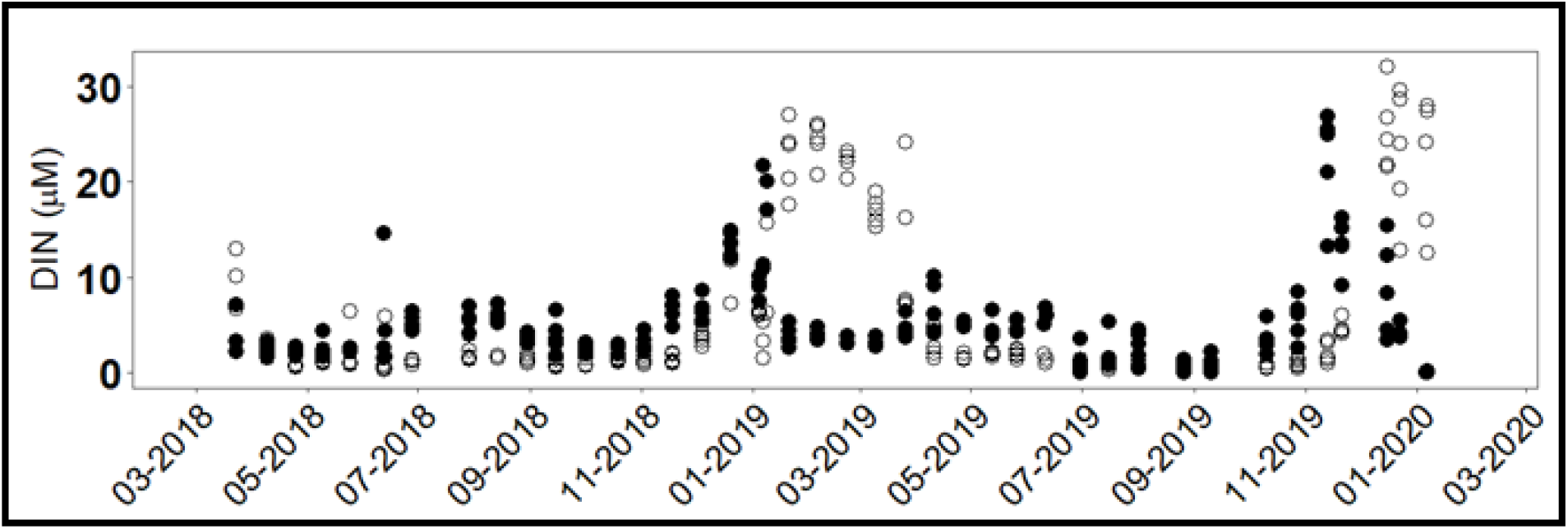
Epilimnetic ammonium (NH_4_^+^) (filled circles) and nitrate (NO_3_^-^) (open circles) at stations B, E, F, P and R.

### 3.4 Nutrient limitation

To assess how these changes in accumulation of hypolimnetic DIN and release with mixing have altered the productivity of the lake, we evaluated nutrient limitation by comparing epilimnetic DIN:TP (by atom) ratios between each sampling period and conducting a nutrient limitation assay. In the historic period, the ratio of DIN:TP in the epilimnion fluctuated between 1.9 and 65.6 (Figure 5a). In contemporary sampling we observed a greater annual range of DIN:TP (1.0 to 564.3) in the photic zone of Lake Yojoa with a pronounced increase in DIN:TP directly following November mixing (Figure 5b).

**Figure 5.**
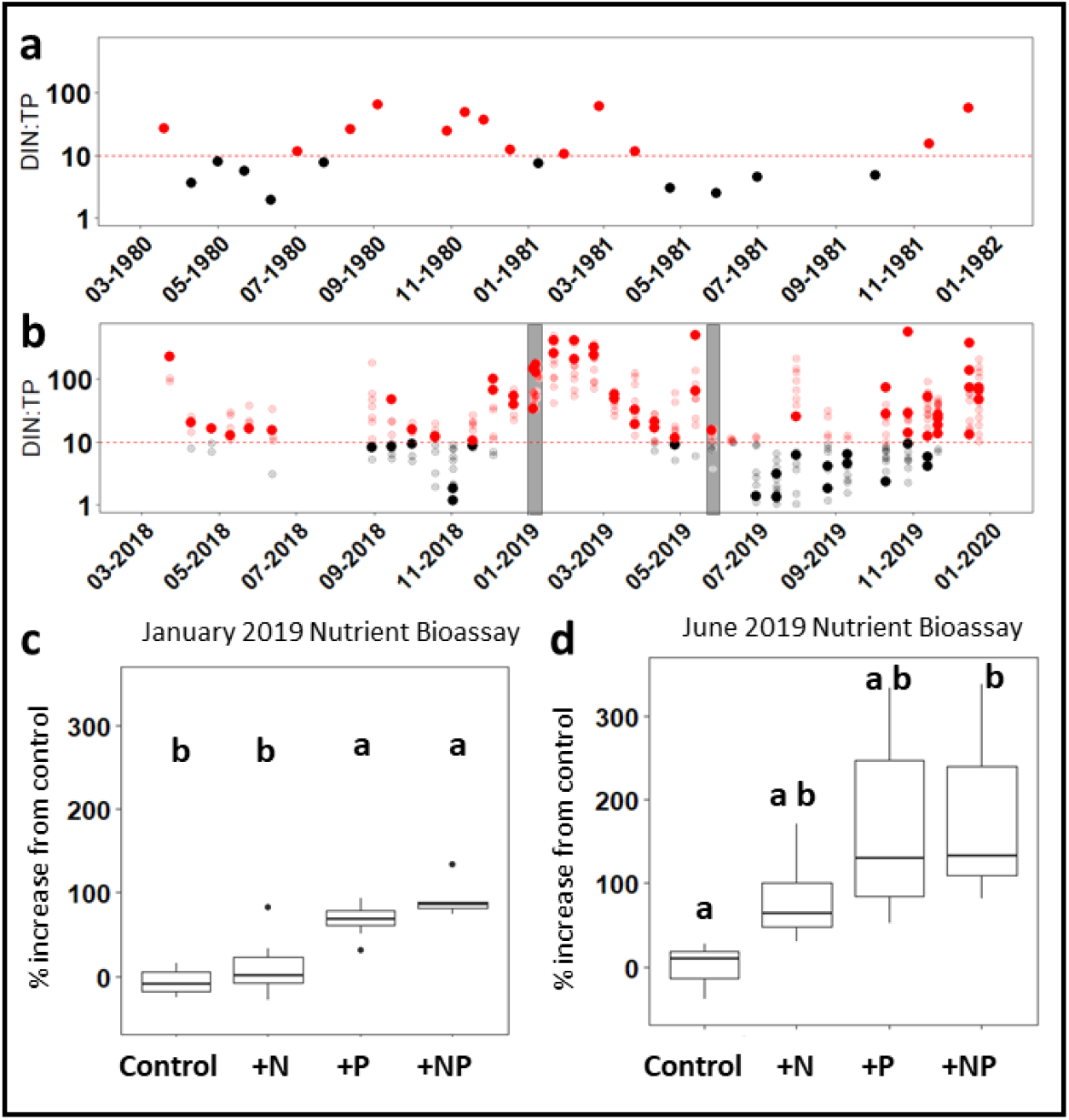
**a)** Historic epilimnetic DIN:TP at location E **b)** contemporary DIN:TP at all locations (B, F, R, P and E bolded). Red indicates likelihood of P limitation above the red dashed line at DIN:TP = 10. P limitation threshold suggested by Wetzel (2001). Grey bars indicate timing of nutrient enrichment bioassays **c)** Nutrient enrichment bioassay from location E conducted January 2019. **d)** Nutrient enrichment bioassay from location E conducted June 2019.

We then directly tested the current state of macronutrient limitation in Lake Yojoa with two nutrient enrichment bioassays, one conducted in January when the water column was mixed, and one conducted in June when the water column was stratified. In January, additions of N alone did not yield a significant increase in algal abundance relative to the control (*p*=0.52, df=32). However, algal abundance was significantly greater in the +P and +NP treatment groups than in both the control and +N treatments (*p*<0.001, df= 32, Figure 5c). The +P and +NP groups were not significantly different from each other (*p*=0.18, df=32). In June, all treatments had a greater percent increase from the control than in January (Figure 5d). However, only treatments that received both N and P (+NP) were significantly different from the control group (*p*=0.054, df=17).

### 3.5 Potential drivers

Whereas this study was not designed to quantify the relative contributions of various point and non-point sources of nutrients to Lake Yojoa, we were able to assess several putative drivers which were most likely to be responsible for the pronounced change in nutrient dynamics. To evaluate the sources of nutrients to contemporary Lake Yojoa we assessed intra-annual delivery of allochthonous (external) nutrients from the watershed using daily cumulative precipitation as a proxy for nutrient loading from the watershed. We compared daily cumulative precipitation in 2019 to epilimnetic DIN and TP concentrations in the pelagic zone of the lake (B, E, F P, and R) to assess the relationship between peak rainfall events and pelagic nutrient concentrations. We found no corresponding increase in epilimnetic nutrient concentrations concomitant with large precipitation events for any of the five principal sampling stations (Figure 6).

**Figure 6.**
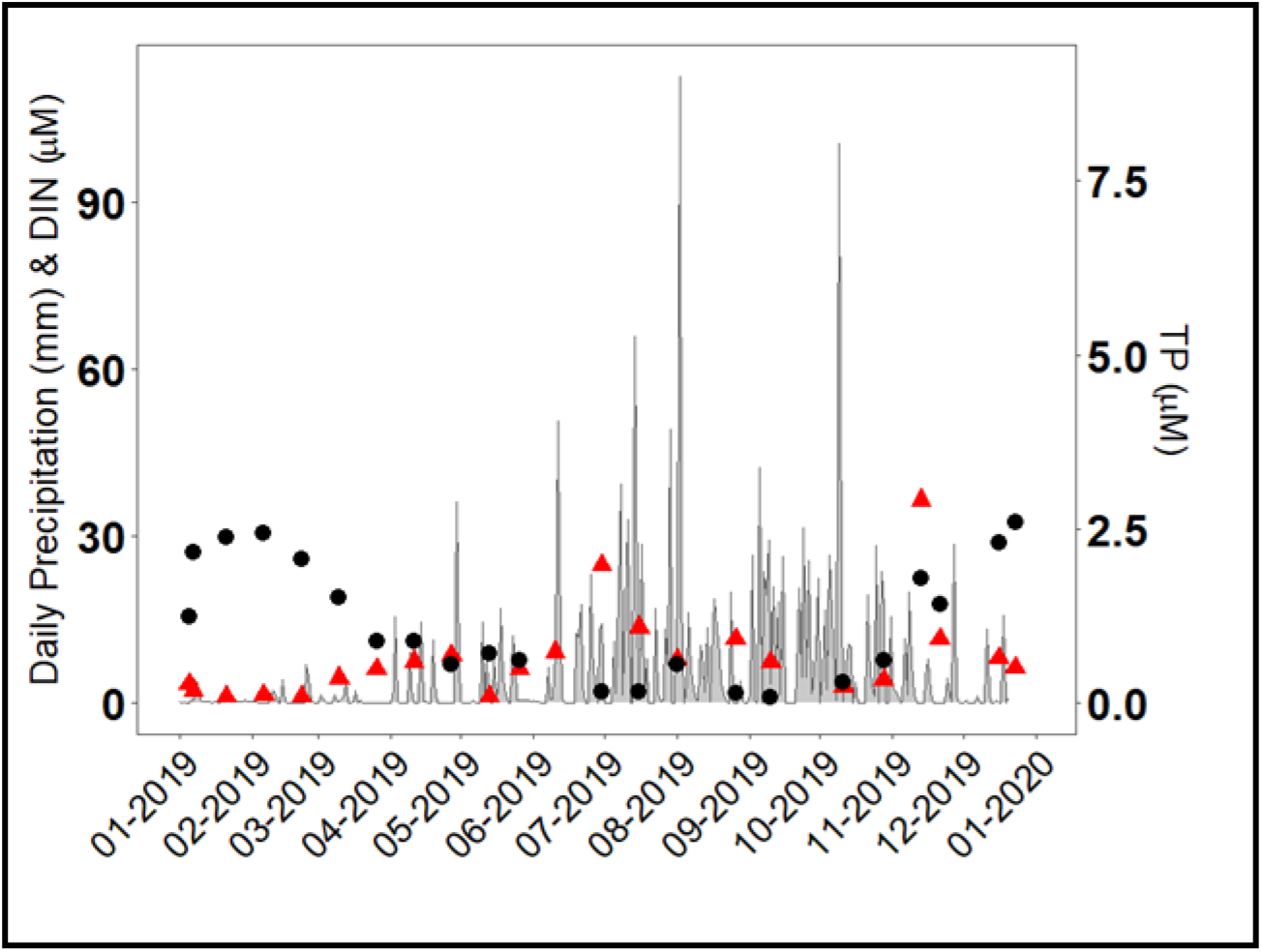
Daily precipitation (primary axis) and mean epilimnetic DIN (black circle) and TP (red triangles) at locations B, E, F, P and R (secondary axis).

We also evaluated the spatial distribution of contemporary hypolimnetic nutrient values during June 2018 and 2019 sampling campaigns. We hypothesized that if nutrient load from tributaries was a major contributor to hypolimnetic nutrient accumulation, hypolimnetic nutrient concentrations would be greatest at sampling stations nearest tributaries (locations B, C, M, N, and O). For the June sampling campaigns, hypolimnetic NH_4_^+^ concentrations in Lake Yojoa ranged from 6.3 to 67.4 μM. However, mean hypolimnetic NH_4_^+^ concentrations at stations nearest the tributaries (locations B, C, M, N, and O mean = 39.03 ± 26.02 μM) were nearly 17% less than the total hypolimnetic mean (47.38 ± 18.22 μM, Figure 7a, Supplementary Table 2) whereas hypolimnetic NH_4_^+^ concentrations at stations in the deepest portion of the lake, Q, P, and R (55.79 ± 11.42 μM) were, on average, approximately 17% greater than the total hypolimnetic mean. In contrast, hypolimnetic NO_3_^-^ concentrations were uniformly low (< 3 μM) at all stations of the lake (Figure 7b, Supplementary Table 2) suggesting that NH_4_^+^ is the dominant form of reactive N in the hypolimnion of Lake Yojoa. Hypolimnetic TP concentrations ranged from 0.6 μM to 3.2 μM. TP concentrations nearest the tributaries (mean= 1.39 ± 0.74 μM) were approximately 37% less than the total hypolimnetic mean (2.21 ± 1.04 μM) and stations in the center of the lake (Q, P and R) had approximately 63% greater TP concentrations (3.58 ± 0.21 μM) than the total hypolimnetic mean (Figure 7c, Supplementary Table 2). Secchi depth was statistically indistinguishable across regions (*p* >0.5) with total lake mean (2.79 ± 0.30 m), near-tributary mean (2.68 ± 0.32 m), and mean of stations Q, P and R (2.85 ± 0.30 m) all within the standard deviation of Secchi depth for the entire lake (Figure 7d, Supplementary Table 2).

**Figure 7.**
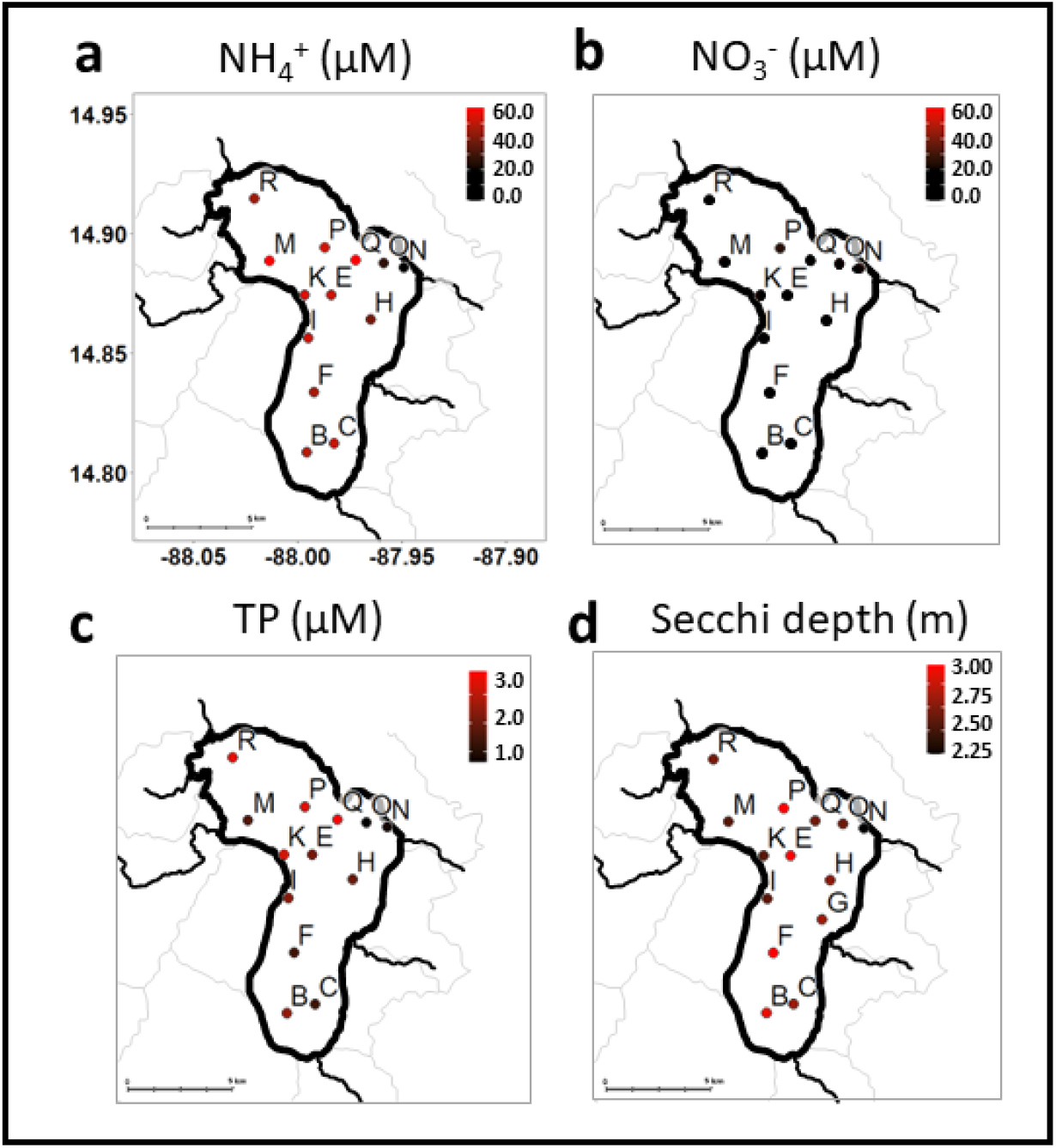
**a)** Mean hypolimnetic NH_4_^+^ **b)** NO_3_^-^ **c)** TP and **d)** Secchi depth in June (mean 2018 and 2019) during contemporary sampling. Values provided in Supplementary Table 2.

## 4. Discussion

Comparison of historic and contemporary periods showed that over the past forty years Lake Yojoa has transitioned from an oligotrophic system with a distinct clear water phase (August to March) to a mesotrophic system with low water clarity and the absence of any pronounced seasonal differences in water clarity. This change in trophic state was accompanied by accumulation of hypolimnetic nutrients (NH_4_^+^ and to a lesser extent TP) during the stratified period and a release of nutrients to the epilimnion during mixing (usually in November). Delivery of NH_4_^+^ from the hypolimnion to the photic zone during the dry season maintained Lake Yojoa’s mesotrophic state by alleviating previous N limitation (replacing phytoplankton N and P colimitation during the stratified water column months with P limitation, Figure 5), allowing rapid uptake of P (also released from the hypolimnion) and preventing the development of the clear water phase that was present during the historic period. To explore drivers and consequences of these changes in the context of contemporary Lake Yojoa, as well as a broader understanding of similar monomictic tropical lakes, in this discussion we focus on 1) potential sources of nutrients to Lake Yojoa, 2) implications for changing nutrient dynamics in other monomictic tropical lake systems, and 3) areas for further research.

### 4.1 Sources of nutrients to Lake Yojoa

The lake and its watershed maintain several livelihood generating activities including approximately 55 lakeside restaurants, agricultural activities (e.g., cultivation of yuca, sugar cane, coffee, and pineapple), subsistence fishing, ranching, and industrial aquaculture (Studer 2007). Transport of reactive nutrients derived from such activities within the watershed undoubtedly influence the biogeochemistry of Lake Yojoa. However, we did not see expected correlations between indicators of watershed loading and nutrient concentrations in the lake, as has been observed in other watersheds (Pandey and Verma 2004). Peak epilimnetic nutrient concentrations did not coincide with the rainy season when the majority of watershed discharge is delivered to the lake (Figure 6). There are a few possible explanations for this observed asynchronicity. First, because the tributaries originate at higher elevations than the lake itself, the disconnect between epilimnetic nutrient concentrations and rain events could be explained by the subduction of cooler tributary waters and the formation of profile-bound density currents which could transport nutrients away from the littoral zone throughout the lake’s hypolimnion, as has been observed in other systems (Talling and Lemoalle 1998). We hypothesized that increased hypolimnetic nutrient concentrations near the mouths of each tributary would suggest that loading from the watershed may be driving the accumulation of hypolimnetic nutrients. However, hypolimnetic sampling stations nearest the littoral zone near the mouths of tributaries were depleted rather than enriched in NH_4_^+^ and TP compared to sampling stations in the hypolimnion of the pelagic zone of the lake (Figure 7a and c). Second, the asynchronicity between watershed loading (as estimated by precipitation) and epilimnetic nutrient concentrations may be explained by nutrient assimilation driven by co-limitation, as suggested by the June nutrient bioassays, leaving reactive nutrients available for only a short period. High rates of nutrient allocation to biomass, not only in phytoplankton but also in aquatic macrophytes has been previously observed in tropical ecosystems (Esteves 1982). However, Secchi depth was not significantly lower nearest the tributaries as would be expected if reactive nutrients were driving spatial differences in algal abundance or if watershed inputs during the rainy season were primarily in the form of suspended sediments and particulate C and N, as observed in other tropical watersheds (McDowell and Asbury 1994). Therefore, we propose that the lack of spatial or temporal correlation between the watershed inputs and water column nutrient concentrations are consistent with the watershed not being the main driver of nutrient changes in Lake Yojoa.

Within the lake are two small “artisanal” net-pen Tilapia farms, one in the central region and another in the southern region of the lake. There is also a large “industrial” net-pen Tilapia operation in the north-central region of the lake (positioned nearest stations P, Q, and R, Figure 1). Net-pen aquaculture cannot treat effluent, causing nutrient rich fish waste (both urea and fecal pellets) to directly mix with the surrounding ecosystem. In other lakes, net-pen aquaculture has been shown to increase the trophic state and contribute to sediment nutrient concentrations (Troell and Berg 1997). Using data shared by the operator of the net-pen aquaculture farm (Regal Springs) for 2011 and 2013, we estimated annual nutrient loading from the aquaculture operation was between 353 to 395 metric tonnes of N and 44 to 55 metric tonnes of P year^-1^ (*pers. comm*. Regal Springs). In addition to the loading of dissolved nutrients, carbon (C) loading (particulate and dissolved) from the Tilapia farm may also be significant given the over 9000 tonnes of feed supplied to the Tilapia pens (based on 2011 and 2013 values), with loading estimates from previous studies on net-pen Tilapia farms ranging from 81 to 91% of C in feed ultimately transferring to the surrounding environment (Gondwe et al. 2011). These estimates suggest the industrial aquaculture operation in Lake Yojoa represents a significant source of C, N, and P to the lake and is a source that was not present during the historic sampling campaign. The importance of the nutrients derived from net-pen aquaculture to the observed changes in Lake Yojoa are also supported by our observations of enriched hypolimnetic NH_4_^+^ and TP at stations nearest the aquaculture operation (P, Q and R) relative to the other stations in the lake, including those nearest the tributary mouths (Figure 7a, c). Despite achieving the highest standards of industry sustainability, including certifications from the Global Aquaculture Alliance/Best Aquaculture Practice (BAP) and the Aquaculture Stewardship Council (ASC), it is evident that the current aquaculture operation is likely introducing unsustainable quantities of nutrients into Lake Yojoa that are altering the state of the ecosystem.

Another potential source of N loading to Lake Yojoa is atmospheric N deposition. While direct measurements of N deposition were unavailable for Lake Yojoa, estimates for the region range from 4.81 – 6.21 mmol –N m^-2^ year^-1^ (Dentener 2006). Given the surface area of Lake Yojoa (88 km^2^, Supplementary Figure 1), this would represent an annual deposition of between 4.23 × 10^5^ and 5.46 × 10^5^ mols per year, approximately two orders of magnitude less than inputs of reactive N derived from aquaculture.

### 4.2 Implications of ongoing change in Lake Yojoa

Lake Yojoa is a monomictic lake that mixes each November (Figure 2a). In 1979-1983 stratification developed in late January or February with the hypolimnion becoming anoxic by the end of March and the thermocline rising to 13-15 m by the beginning of the rainy season (typically in May) and mixing occurring in November (Vaux and Goldman 1984). We observed the same timing of mixus and stratification in contemporary Lake Yojoa but with a slightly shallower thermocline (11-12m) (Figure 2a). Therefore, the contemporary stratification conditions which allow for hypolimnetic nutrient accumulation existed in the 1980s, though in the 1980s both epilimnetic and hypolimnetic temperatures were slightly less (∼ 3 ºC) than what is observed today (Figure 2b). In addition to the predicted seasonal mixing in November, during the contemporary sampling campaign, we also observed infrequent, ephemeral partial mixing events (i.e., only at some stations or not complete in depth with stratification quickly re-establishing) concurrent with large wind events and/or cold fronts (Figure 2a). Similar partial mixing events during the stratified season were also noted in the historic data, as evidenced by anomalies in oxygen and temperature profiles (Vaux and Goldman 1984). However, these aseasonal mixing events have new ramifications under conditions of elevated levels of hypolimnetic nutrients in contemporary Lake Yojoa. Mixing events, during otherwise stratified months, introduce hypolimnetic nutrients to surface waters during peak annual temperatures. This pulse of nutrients can result in rapid increases in algal biomass (i.e., algal blooms). One such bloom occurred on June 9, 2020, and garnered national attention (La Tribuna, 2020). With increased intensity of weather events predicted with climate change (Knutson et al. 2010), these aseasonal mixing events may become more frequent, with the potential for a concurrent increase in frequency of algal blooms during the months where the lake is normally stratified.

The accumulation of hypolimnetic NH_4_^+^ and TP and subsequent release into the photic zone during the dry season when watershed inputs are low is a relatively recent (<40 years) development in Lake Yojoa. However, we hypothesize that a similar pattern of nutrient delivery from the hypolimnion to the photic zone during the dry season may be driving changes in trophic state of other monomictic tropical lakes. For example, annual mixing in Lake Titicaca (Peru-Bolivia) introduced large quantities of nutrients to the euphotic zone, temporarily alleviating N and P limitations (Vincent et al. 1984). Similarly, in Valle de Bravo Reservoir (Mexico), N limitation during stratification was replaced by P limitation following mixing due to an increase in the DIN:TP ratio (Merino-Ibarra et al. 2008). These findings in other monomictic tropical systems are consistent with the results from the nutrient bioassays and the seasonal shifts in DIN:TP we observed in this study (Figure 5).

### 4.3 Other controls on lake N dynamics

To better understand the observed changes of hypolimnetic nutrient accumulation, we estimated changes in the rate of net hypolimnetic NH_4_^+^ and TP accumulation from January, when the water column was mixed, to July 1st, near peak stratification, between contemporary and historic periods. In the early 1980s, NH_4_^+^ accumulated in the hypolimnion at an estimated rate of 0.04 μM day^-1^. Today, we estimate an accumulation rate of 0.33 μM day^-1^ over the same interval, a nearly tenfold increase in the rate of NH_4_^+^ accumulation in the hypolimnion between the historic and contemporary periods. We estimate that TP accumulated at a rate of 0.0005 μM day^- 1^ in the historic period compared to 0.002 μM day^-1^ in the contemporary. Our calculations here suggest that this order of magnitude difference in N and P loading to the hypolimnion is likely primarily driven by the industrial aquaculture operation that was established in Lake Yojoa in 1997.

Given the warm, anoxic hypolimnion of Lake Yojoa, it is also likely that unique anaerobic microbial pathways may also contribute to the accumulation of NH_4_^+^ in the hypolimnion. Changes in N-cycling pathways that contribute (and remove) NH_4_^+^ may be an important aspect of the understanding how hypolimnetic processes contribute to Lake Yojoa’s changing trophic state. There are seven principal biogeochemical pathways that have the potential to influence the pool of NH_4_^+^ in of Lake Yojoa: N fixation, nitrification, uptake, mineralization, denitrification, anaerobic ammonium oxidation (anammox) and dissimilatory nitrate reduction to ammonium (DNRA). Of these, only uptake, mineralization, denitrification, anammox, and DNRA, are likely to be active in the aphotic, anoxic hypolimnion of Lake Yojoa. Thus, understanding controls and constraints on each of these pathways may provide further insight into the changing biogeochemistry of monomictic tropical lakes. For example, the large inputs of organic N from the fish farm as well as the warm temperatures present in the hypolimnion of Lake Yojoa likely result in accelerated rates of mineralization (Amado et al. 2013). In addition, concentrated organic matter loading below aquaculture operations has been shown to increase DNRA rates up to seven-fold of ambient conditions (Christensen et al. 2000) further exacerbating the accumulation of ammonium. Under conditions of low NO_3_^-^, as observed in Lake Yojoa’s hypolimnion (Figure 7b) and high concentrations of labile C, DNRA is favored over complete denitrification, the dominant loss pathway for reactive N in most lake ecosystems (Nizzoli et al. 2010). Given the large inputs of C that are likely associated with the industrial Tilapia operation in the deepest basin of the lake, conditions that favor DNRA are likely present in the hypolimnion of Lake Yojoa. Conditions that result in the selection for DNRA over denitrification would further promote the accumulation of NH_4_^+^, whereas historic conditions (e.g., lower autochthonous N and organic C inputs) may have favored denitrification and prevented the accumulation of hypolimnetic NH_4_^+^. Additional studies on controls over microbially mediated N transformation pathways in the warm, anoxic hypolimnions may be a key tool to better understand ecosystem scale change in tropical lakes.

## 5. Conclusions

The results from this study along with similar observations in other monomictic tropical lakes suggest that changes in hypolimnetic nutrient accumulation and release to the photic zone coincident with mixus are driving changes in the trophic state of monomictic tropical lake ecosystems (Vincent et al. 1984, Merino-Ibarra et al. 2008). In Lake Yojoa, the presence of large-scale aquaculture is likely driving the loading of N and P to the hypolimnion but in other lakes the source of N or P may be different. In monomictic tropical lakes, the interaction of the physical structure of the water column and nutrient loading places hypolimnetic processes at the intersection of local drivers, such as nutrient loading from aquaculture, and global drivers, such as warming waters and increasing storm intensity associated with climate change (Knutson et al. 2010). Our results suggest that one key to improving the current understanding of tropical monomictic lake ecosystems may be defining the controls on hypolimnetic biogeochemical pathways. Comparisons between epilimnetic and hypolimnetic nutrient concentrations may therefore be an important indicator of trophic state threshold in monomictic lake ecosystems and provide a useful tool to assess the impact of contemporary uses and stressors on the overall trophic state of these societally important ecosystems.

## Acknowledgements

We would like to thank the entire staff of AMUPROLAGO for their efforts in this collaboration, in particular Juan Carlos Sorto for sample collection. Regal Springs Inc. provided N and P loading estimates from their aquaculture operation. We thank Dr. Charles Goldman and Dr. Peter Vaux for sharing the original, hand-written data from the historic study. This project benefitted greatly from the support of Casey Barby both in the field and in the lab. This manuscript was greatly improved by valuable feedback from Dr. John Melack, Dr. André Amado and two anonymous reviewers. Funding was provided by the National Science Foundation [DEB # 2120441, 2021], Warner College of Natural Resources, and Department of Ecosystem Science and Sustainability at Colorado State University (CSU) with additional support from the Graduate Degree Program in Ecology and the Natural Resource Ecology Lab at CSU. Additional travel support was provided by two airfare vouchers from United Airlines. We have no conflicts of interest to declare.

