## Supplemental Table 1 for "The interaction of physical structure and nutrient loading drives ecosystem change in a large tropical lake over 40 years"

**Supplementary Table 1.** Physical characteristics of Lake Yojoa 1) Vaux and Goldman (1984) 2) Romero and Pineda (2007).

| | |
|---|---|
| Lake Area ($km^2$) | 88[1] |
| Lake Volume ($km^3$) | 1.4[2] |
| Elevation (m above sea level) | 638[1] |
| Catchment area ($km^2$) | 337[1] |
| Mean depth (m) | 14.6[2] |
| Maximum depth (m) | 27.3[2] |
| 1) Vaux and Goldman (1984) 2) Romero and Pineda (2007). | |
