## Supplemental Table 2 for "The interaction of physical structure and nutrient loading drives ecosystem change in a large tropical lake over 40 years"

**Supplementary Table 2.** Mean hypolimnetic nutrient values for June 2018 and 2019 with corresponding Secchi depth.

| | $NH_4^+$ (µM) | $NO_3^-$ (µM) | TP (µM) | Secchi depth (m) |
|---|---|---|---|---|
| A | *Shallow, no hypolimnion* | | | |
| B | 51.13 | 1.22 | 2.48 | 3.1 |
| C | 53.88 | 0.97 | 1.40 | 2.8 |
| D | *Shallow, no hypolimnion* | | | |
| E | 55.47 | 0.90 | 2.05 | 3.1 |
| F | 46.18 | 0.83 | 1.40 | 3.2 |
| G | *Shallow, no hypolimnion* | | | 2.8 |
| H | 35.15 | 0.67 | 1.95 | 2.7 |
| I | 57.24 | 1.09 | 2.35 | 2.6 |
| J | *Shallow, no hypolimnion* | | | |
| K | 59.36 | 1.50 | 3.22 | 2.5 |
| L | *Shallow, no hypolimnion* | | | |
| M | 66.95 | 2.47 | 1.68 | 2.5 |
| N | 6.31 | 1.61 | 0.79 | 2.2 |
| O | 16.88 | 2.99 | 0.62 | 2.6 |
| P | 55.47 | 1.00 | 3.42 | 3.2 |
| Q | 67.37 | 1.11 | 3.82 | 2.6 |
| R | 44.54 | 0.93 | 3.49 | 2.7 |
