## Appendix A for "The interaction of physical structure and nutrient loading drives ecosystem change in a large tropical lake over 40 years"

The relationship between Secchi depth and algal biomass in contemporary Lake Yojoa across all sampled months of the contemporary sampling period is statistically significant ($p < 0.05$) if not particularly strong ($R^2 = 0.49$) (Figure 1). The lack of a stronger relationship is likely attributable to some of the well-documented challenges in using chlorophyll a as a direct proxy for algal biomass[1] coupled with high variability in algal flocculation we observed in Lake Yojoa (resulting in variability in estimates of volumetric chlorophyll concentrations for any given sampling event). Looking only at the months when the water column is mixed and when flocculation is uncommon (November-February), the relationship between Secchi depth and algal abundance is much stronger ($p<0.000005$, $R^2 = 0.86$) (Figure 2). Thus, in Lake Yojoa the vast majority of variance in Secchi depth can be explained by shifts in algal abundance.

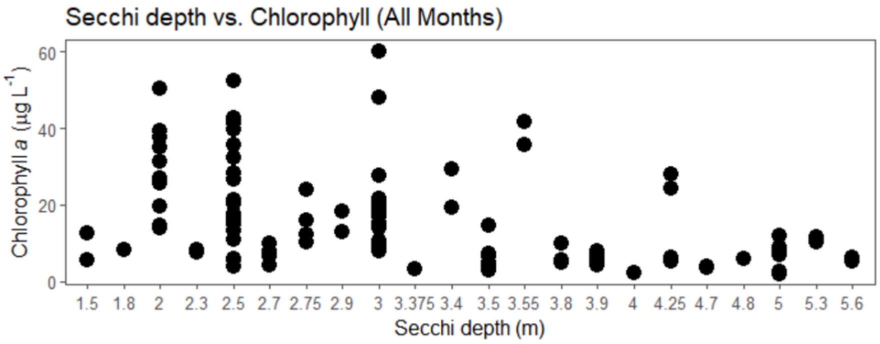


**Figure 1.** Secchi depth vs algal biomass, as estimated by chlorophyll a concentration, for all study months.

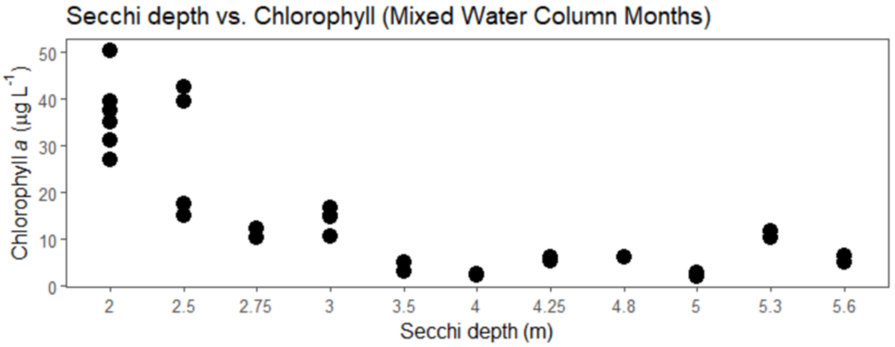


**Figure 2.** Secchi depth vs algal biomass, as estimated by chlorophyll a concentration, for months when the water column of Lake Yojoa is mixed (November-February).

---

[1] Ramaraj, R et al. 2013. Chlorophyll is not accurate measurement for algal biomass. Chiang Mai J. Sci. 40:1-9.
